# Susceptibility to Neutralization by Broadly Neutralizing Antibodies Correlates with Infected Cell Binding for a Panel of Clade B HIV Reactivated from Latent Reservoirs

**DOI:** 10.1101/330894

**Authors:** Yanqin Ren, Maria Korom, Ronald Truong, Dora Chan, Szu-han Huang, Colin C. Kovacs, Erika Benko, Jeffrey T. Safrit, John Lee, Hermes Garbán, Richard Apps, Harris Goldstein, Rebecca M. Lynch, R.Brad Jones

## Abstract

Efforts to HIV cure are obstructed by reservoirs of latently infected CD4^+^ T-cells that can re-establish viremia. Broadly neutralizing HIV-specific antibodies (bNAbs), defined by unusually high neutralization breadths against globally diverse viruses, may contribute to the elimination of these reservoirs by binding to reactivated cells, targeting them for immune clearance. However, the relationship between neutralization of reservoir isolates and binding to corresponding infected primary CD4^+^ T-cells has not been determined. Thus, the extent to which neutralization breadths and potencies can be used to infer the corresponding parameters of infected-cell binding is currently unknown. We assessed the breadths and potencies of bNAbs against 36 viruses reactivated from peripheral blood CD4^+^ T-cells of ARV-treated HIV-infected individuals, using paired neutralization and infected-cell binding assays. Single antibody breadths ranged from 0–64% for neutralization (IC_80_≤10μg/ml) and 0–89% for binding, with two-antibody combinations reaching 0-83% and 50-100%, respectively. Infected-cell binding correlated with virus neutralization for 10 out of 14 antibodies (e.g. 3BNC117, r=0.87, p<0.0001). Heterogeneity was observed, however, with a lack of significant correlations for 2G12, CAP256.VRC26.25, 2F5, and 4E10. Our results provide guidance on the selection of bNAbs for interventional cure studies; both by providing a direct assessment of intra-and inter-individual variability in neutralization and infected cell binding in a novel cohort, and by defining the relationships between these parameters for a panel of bNAbs.

**Importance:** Although anti-retroviral therapies have improved the lives of people who are living with HIV, they do not cure infection. Efforts are being directed towards harnessing the immune system to eliminate the virus that persists, potentially resulting in virus-free remission without medication. HIV-specific antibodies hold promise for such therapies owing to their abilities to both prevent the infection of new cells (neutralization), and also to direct the killing of infected cells. We isolated 36 HIV strains from individuals whose virus was suppressed by medication, and tested 14 different antibodies for neutralization of these viruses and for binding to cells infected with the same viruses (critical for engaging natural killer cells). For both neutralization and infected-cell binding, we observed variation both between individuals, and amongst different viruses within an individual. For most antibodies, neutralization activity correlated with infected cell binding. These data provide guidance on the selection of antibodies for clinical trials.

## Introduction

Modern antiretroviral (ARV) drug regimens effectively suppress HIV replication, but are unable to cure infection. Interruption of ARV therapy thus results in rapid viral rebound and disease progression. A critical aspect of HIV persistence in the context of ARV therapy is the establishment of latent infection in long-lived resting memory CD4^+^ T-cells (1-3). Evidence from *in vitro* latency models supports that these reservoirs can be eliminated by combining latency reversal agents (LRAs), which induce the expression of viral antigens, with enhanced immune effectors; a paradigm referred to as “kick and kill” or, alternatively, as “shock and kill” (4-6). One strategy to harness immune effectors for these strategies is to target reactivated infected cells with HIV-specific antibodies, resulting in the engagement of natural killer (NK) cells, monocytes, and granulocytes which eliminate infected cells through antibody-dependent cell-mediated cytotoxicity (ADCC) and/or antibody-dependent cell-mediated phagocytosis (ADCP) (7-9). For this purpose, it will be crucial for the HIV-specific antibodies to bind to Env protein expressed on the surface of the reactivated latent infected cells. The current study focuses on correlating the susceptibility of neutralization against viral isolates reactivated from patient CD4^+^ T-cells by a panel of HIV-specific broadly neutralizing antibodies (bNAbs) with their capacity to bind to Env expressed by the reactivated latent infected cells, thereby providing guidance on the selection of bNAbs to optimally support the clinical translation of kick and kill strategies.

The antigenic variability of the HIV Envelope protein poses a substantial challenge to the development of both vaccines and immunotherapeutics (10-12). The past 10 years have seen the identification of a growing number of ‘broadly neutralizing antibodies’ (bNAbs), defined as such based on their activity against globally diverse HIV isolates (13-22) [reviewed in (23-26)]. Recent clinical trials have established that passive infusion with bNAbs during chronic HIV infection can temporarily suppress virus replication in individuals whose virus does not escape (27-29), and modestly delay viral rebound during anti-retroviral treatment interruption (30, 31). Additionally, passive immunization with bNAbs has attracted interest as a means of supplying the immune effector component of kick and kill HIV eradication strategies (given that virus has typically escaped from autologous antibody responses). This has led to the initiation of additional preclinical trials, as well as pilot clinical studies aimed at testing the abilities of combinations of bNAbs and latency reversing agents (LRAs) to reduce or eliminate latent HIV reservoirs (e.g. ClinicalTrials.gov NCT03041012, NCT02850016).

Three primary factors argue for the prioritization of bNAbs, versus other types of HIV-specific antibodies, for clinical trials aimed at reducing latent reservoirs through a kick and kill mechanism. First, there is extensive clinical experience with, and safety data on, several bNAbs from their use in passive infusion trials; facilitating their advancement into combination studies with LRAs. Second, the ability to exert the dual activities of neutralizing free virus in addition to mediating ADCC would be favorable for an antibody therapeutic. Third, the antigenic diversity of HIV, both within a given individual’s latent reservoir and at a population level, poses a challenge to the development of curative therapeutics, motivating the prioritization of Abs with broad reactivity. With respect to the latter point, while it stands to reason that an Ab with broad neutralizing activity is likely to exert a similar breadth of infected-cell binding, this cannot be assumed to be the case. Infected cell binding is a prerequisite for, and correlates closely with, ADCC activity (8, 32-34). The conformations of Env on free virions that must be targeted to achieve neutralization may differ from those on infected cells that must be bound to trigger ADCC. For example, binding of Env on an infected cell to CD4 on that same cell (i.e: in *cis*) may both partially occlude the CD4 binding site (CD4bs) and induce gp120 shedding, while exposing CD4-induced (CD4i) epitopes and gp41 stumps (35); thus, antigenically changing the protein on a cell as compared to the virion. Although CD4i antibodies commonly arise during infection (36), and have the potential to mediate ADCC against liganded versions of the Env protein, the addition of sCD4 mimetics has been necessary to increase sensitivity of infected cells to ADCC by these antibodies (37, 38). Furthermore, the possibility exists that viral diversity may differentially affect cell-surface Env versus virion-associated functional Env trimers, potentially in unexpected ways. Thus, broadly neutralizing antibodies present the possibility of infusing multi-functional antibodies that target genetically diverse viruses on epitopes that do not require CD4 binding for epitope exposure; however, broad neutralizing activity may not equate to broad infected-cell binding. Of note, the bNAbs tested in study all share the same IgG1 Fc domain, differing only in their Fab fragments. The current study thus focuses on providing guidance with respect to the selection of the antigen binding Fab fragments of Abs for use in cure strategies. To maximize potency, these Fab fragments may ultimately need to be combined with Fc domains that are designed to maximally engage ADCC effectors (39).

A limited number of studies have thus far assessed the breadths of infected-cell binding and/or ADCC activity by bNAbs in relation to neutralizing activity, and these have reported somewhat conflicting results. In testing 8 viral isolates reactivated from the latent reservoirs of ARV-treated individuals, Bruel *et al.* reported that a panel of bNAbs (including 3BNC117) could eliminate HIV infected cells by mediating ADCC (8), and that their breadth of virus recognition was higher than with non-neutralizing antibodies (32). In contrast, Mujib *et al*. reported a lack of infected-cell binding and ADCC activity by 3BNC117 against a multi-clade panel of HIV (40), suggesting a lack of correspondence with its breadth of neutralizing activity (15). Although this relationship has been explored indirectly, to our knowledge, only one study has directly compared infected cell binding or ADCC of bNAbs versus neutralizing activity across different viral isolates. This study showed a correlation between these functions, but was limited to the use of two viral isolates of HIV (NL4-3 and JR-FL) and SHIV AD8-EO (34). We therefore perceived a need to define the relationship between neutralization and infected cell binding of clinically relevant bNAbs to HIV produced by reactivated latent infected CD4^+^ T cells.

In the current study we assessed, in parallel, virus neutralization and infected primary CD4^+^ T-cell binding of bNAbs against a panel of 36 viruses that were reactivated from the latent reservoirs of 8 ARV-treated individuals by quantitative viral outgrowth assays (QVOA) (41) (see schematic, **Fig 1**). We defined the intra-and inter-patient breadths and potencies of both neutralization and infected cell binding activity of these bNAbs against reactivated reservoir viruses from a geographically localized population of clade B infected individuals. For all bNAbs that demonstrated appreciable neutralizing activity, this correlated closely with infected cell binding. This represents the most comprehensive study to date using a large panel of bNAbs, which target a range of different epitopes but share the same IgG1 Fc domain, against a panel of *ex vivo* reservoir reactivated viruses to quantify both neutralization and binding to infected cells.

**Figure 1.**
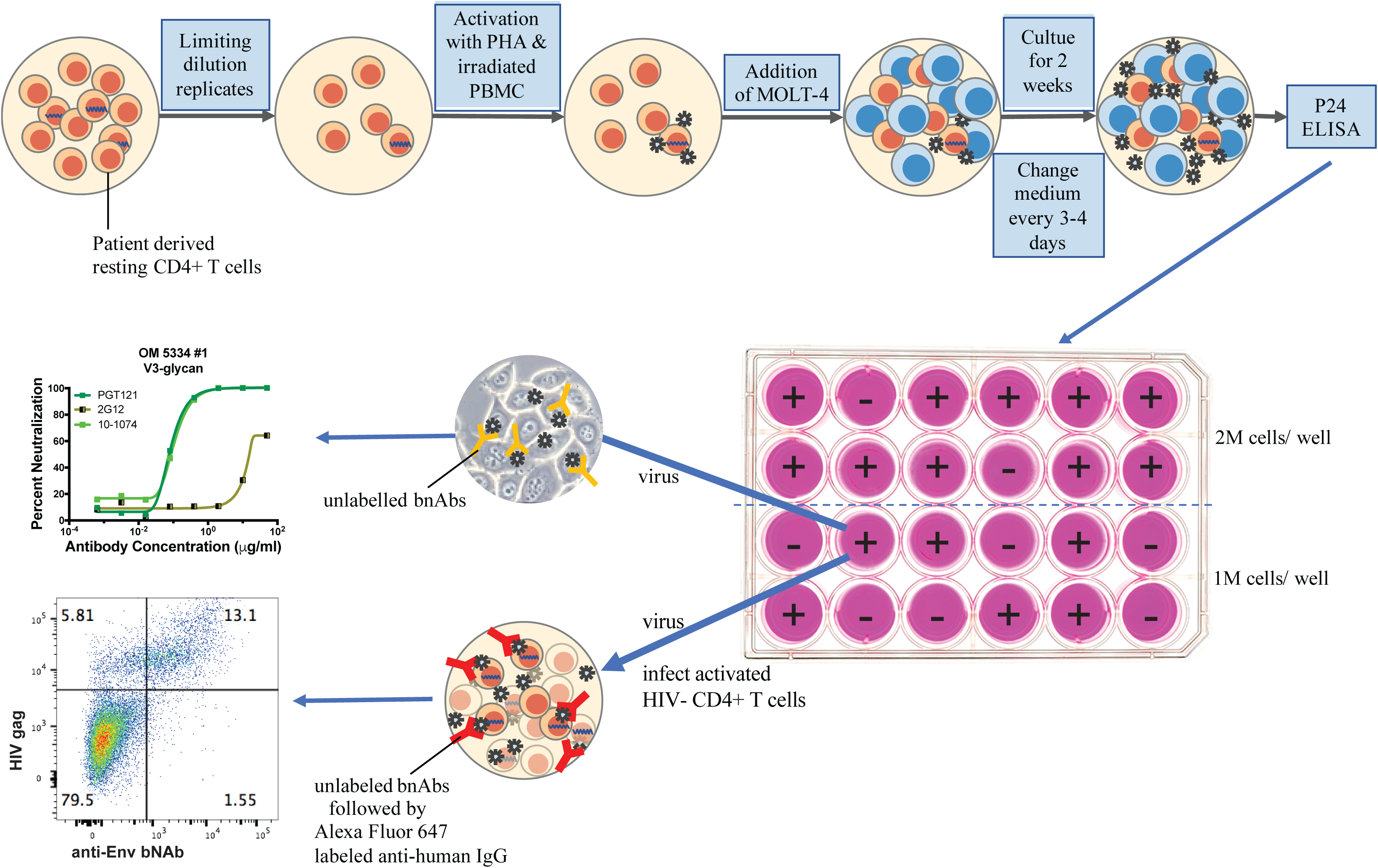
Schematic for paired assessment of virus neutralization and infected cell binding with reactivated reservoir viruses. Quantitative Viral Outgrowth Assays (QVOA) were performed using CD4^+^ T-cells from ARV-suppressed study participants. Virus was isolated from HIV-p24^+^ wells at a dilution where < 50% of wells were positive. A portion of the supernatants from each of these wells was used directly to assess virus neutralization using a TZM-bl assay. Another portion was used to infect activated primary CD4^+^ T-cells. Binding of bNAbs to these infected cells was assessed by flow cytometry, co-staining with CD3, CD4 and HIV-Gag to identify infected cells.

## Results

### Virus Neutralization Profiles of bNAbs and bNAb Combinations Against Reactivated Reservoir Viruses

To test the ability of bNAbs to neutralize reservoir virus, we obtained a panel of 14 bNAbs that are currently being developed for clinical use in humans and categorized these by their targeted epitope (**See Methods**). We measured the neutralizing activities of these bNAbs against 36 viral isolates that had been reactivated from the latent reservoirs of 8 individuals from limiting dilution quantitative viral outgrowth assays (QVOA) (**Fig 1 and 2A**). The V3-glycan-specific bNAbs PGT121 and 10-1074 and the V1V2-specific bNAb PG9 exhibited potent but relatively narrow activity, exhibiting detectable neutralization (IC_50_ < 50 μg/ml) of 53 - 69% of viruses, with geometric mean IC_50_ values ranging from 0.3 – 0.6 μg/ml (**Fig 2B**). In contrast, the CD4 binding site (CD4bs)-specific antibodies VRC01, VRC07-523, N6 and 3BNC117, as well as the MPER-targeting antibody 10E8 exhibited broad activity, with a detectable neutralization 77 - 100% of viruses (IC_50_ < 50 μg/ml), but with substantially higher IC_50_ values (geometric mean IC_50_ between 2.1 – 8.9 μg/ml) (**Fig 2B**). These trends parallel previous reports using pseudovirus assays, which also observed that CD4bs antibodies and 10E8 were generally much broader but less potent than V3-glycan and V1V2 apex antibodies (42, 43). In the current experiment, CAP256.VRC26.25 only neutralized 9 of 36 reactivated reservoir viruses (26%) with a detectable IC_50_ (IC_50_ < 50 μg/ml) (**Fig 2B**). Because CAP256.VRC26.25 has been reported to preferentially neutralize subtype C, and the QVOA viral isolates tested here are all subtype B (**Table 1**), the low neutralization breadth we observed is compatible with published data (22). 4E10 and 2F5 are known to be less broad and potent than more recently published antibodies, so their lack of breadth against these viruses is expected. One exception to the general agreement between our data and those from published pseudovirus panels was for 2G12 which, although not broadly neutralizing against genetically diverse viruses, has been shown to potently neutralizes subtype B viruses in published pseudovirus panels (19, 44), but we observed only weak neutralization in our assays, with only two viruses reaching 80% neutralization (**Fig 2**).

**Figure 2.**
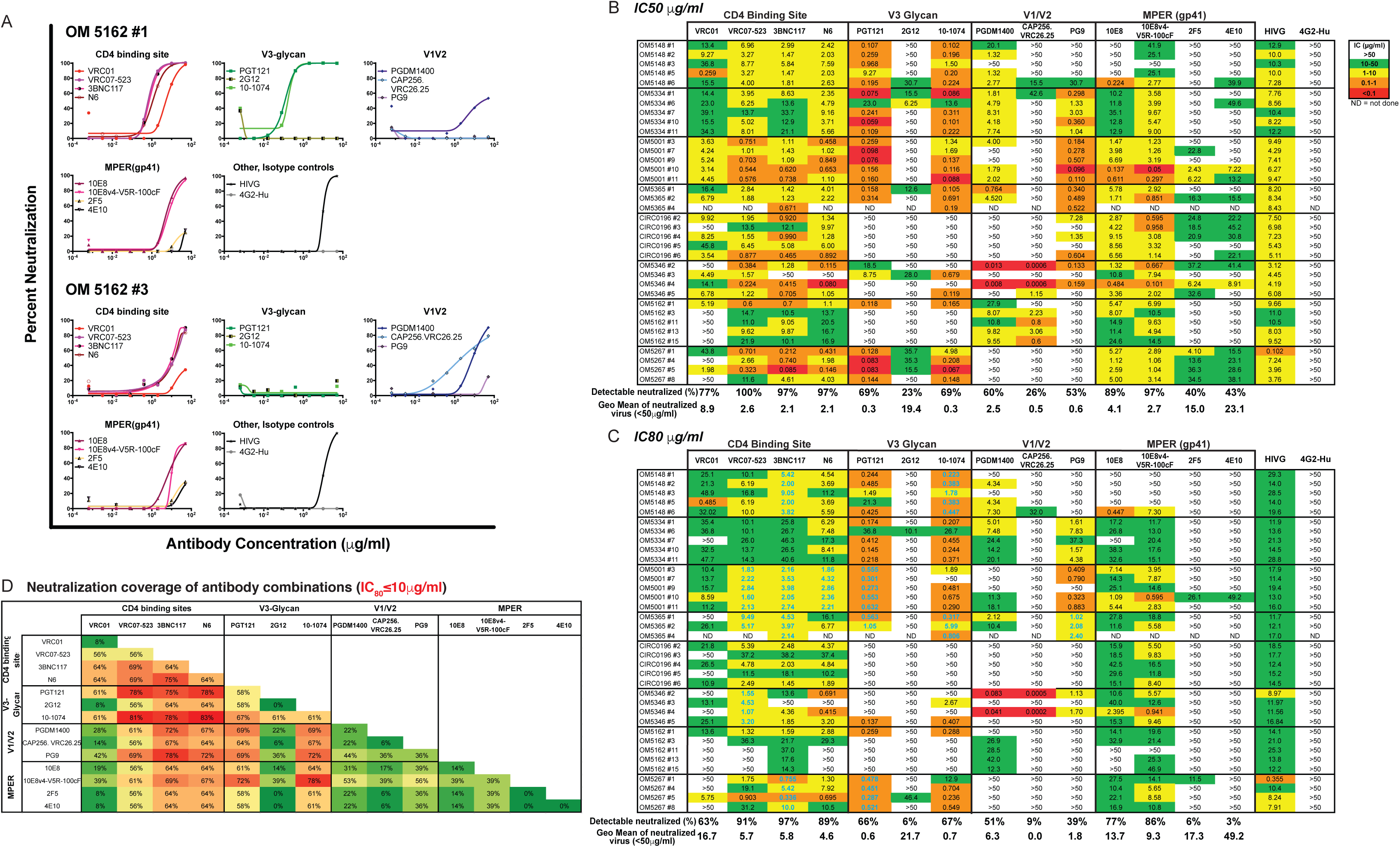
Breadth and potency of neutralization of a panel of bNAbs against reactivated reservoir viruses. (A) Representative neutralization curves against virus isolates #1 & #3 from study participant OM5162. Each graph represents antibodies targeting similar epitopes against one virus, and each curve represents results from one bNAb. (B) The half maximal inhibitory concentration (IC_50_) and (C) IC_80_ are shown in heat-maps. The lower the antibody concentration, the more sensitive the reservoir virus is to a specific bNAb (bNAbs were shown by binding epitope classes; HIV-IG, positive control antibody; 4G2-Hu, negative control antibody). The geometric mean concentration against all 36 (or 35) reservoir viruses tested was calculated. Numbers in Cyan bold are those where a single bNAb provided coverage of each of the viral isolates tested from a given participant. (D) Heat-map showing neutralization coverage of antibody combinations. Shown are the % of viral isolates that were neutralized by at least one antibody in the indicated combinations using an IC_80_ cut off of 10 μg/ml.

**Table 1.**
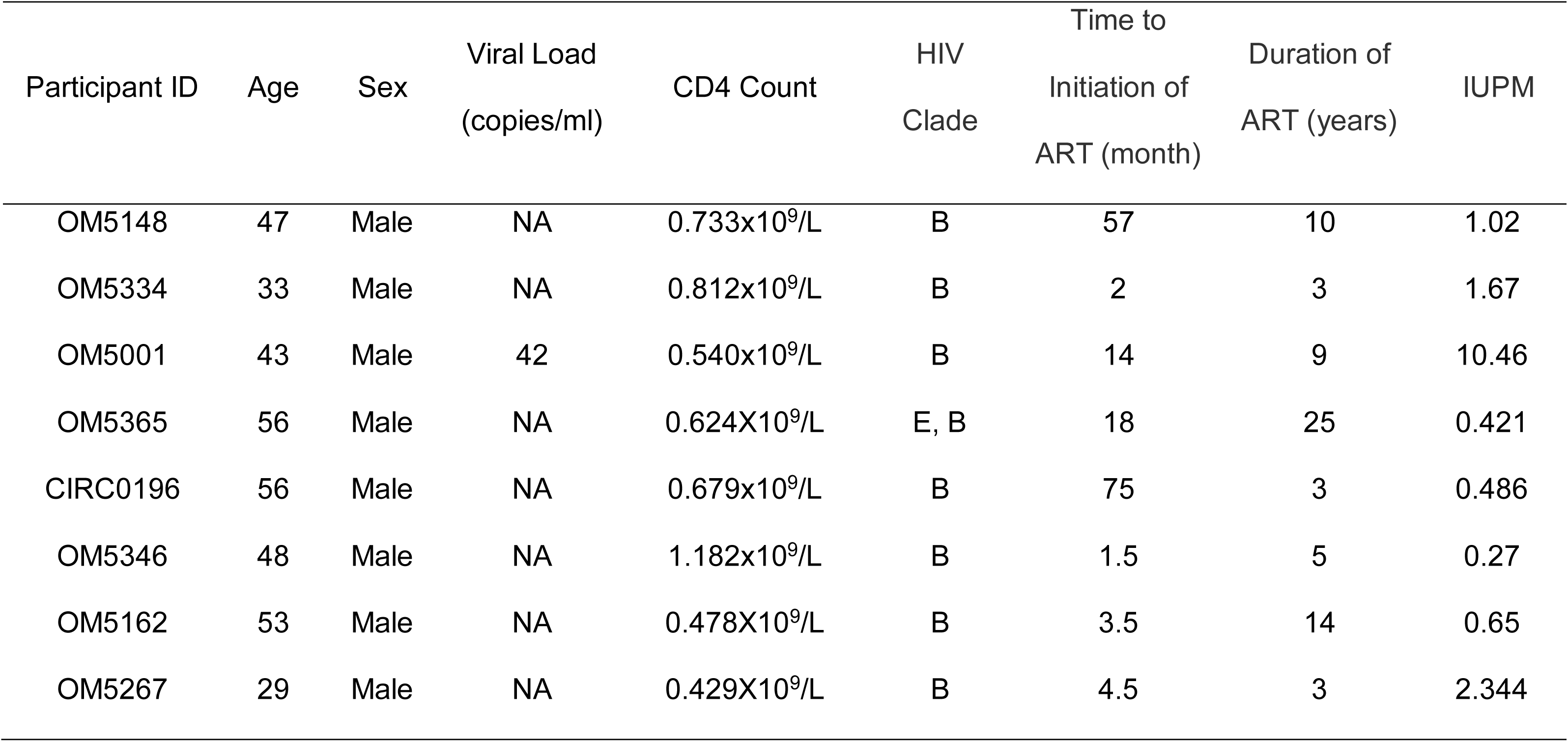
Patient clinical information.

| Participant ID | Age | Sex | Viral Load<br>(copies/ml) | CD4 Count | HIV<br>Clade | Time to<br>Initiation of<br>ART (month) | Duration of<br>ART (years) | IUPM |
| --- | --- | --- | --- | --- | --- | --- | --- | --- |
| OM5148 | 47 | Male | NA | 0.733x10<sup>9</sup>/L | B | 57 | 10 | 1.02 |
| OM5334 | 33 | Male | NA | 0.812x10<sup>9</sup>/L | B | 2 | 3 | 1.67 |
| OM5001 | 43 | Male | 42 | 0.540x10<sup>9</sup>/L | B | 14 | 9 | 10.46 |
| OM5365 | 56 | Male | NA | 0.624X10<sup>9</sup>/L | E, B | 18 | 25 | 0.421 |
| CIRC0196 | 56 | Male | NA | 0.679x10<sup>9</sup>/L | B | 75 | 3 | 0.486 |
| OM5346 | 48 | Male | NA | 1.182x10<sup>9</sup>/L | B | 1.5 | 5 | 0.27 |
| OM5162 | 53 | Male | NA | 0.478X10<sup>9</sup>/L | B | 3.5 | 14 | 0.65 |
| OM5267 | 29 | Male | NA | 0.429X10<sup>9</sup>/L | B | 4.5 | 3 | 2.344 |

We frequently observed high degrees of similarity in neutralization sensitivities within an individual’s viral quasispecies, consistent with genetic relatedness. For example, the five viral isolates from CIRC1096 were all sensitive to neutralization by CD4bs and MPER antibodies, but resistant to V3-glycan and V1V2 antibodies (**Fig 2B & C**). Exceptions to this, however, were not uncommon. For example, for the four QVOA viruses from OM5346, two of these viruses (#2 and #4) were highly sensitive to V1V2 antibodies (PG9, CAP256-VRC26.25, PGDM1400) and resistant to V3-glycan antibodies (PGT121, 10-1074 and 2G12) whereas virus #3 exhibited the opposite sensitivity profile (**Fig 2B & C**). Overall, of the 112 study participant / bNAb combinations (8 participants × 14 bNAbs) there were only 14 cases where a single bNAb provided coverage of each of the viral isolates tested from a given participant (IC_80_ ≤ 10 μg/ml, **Fig 2C in Cyan Bold**).

Given the limitations observed above in the breadths of coverage and potential escape of any single bNAb, it is likely that any clinical intervention would require combinations of multiple bNAbs to be effective. We therefore calculated the summed breadths of all combinations of two of the bNAbs tested in this study. We determined breadth coverage by using an IC_80_ ≤ 10 μg/ml as the cut-off for the geometric mean sensitivity of the quasispecies, based on our previous demonstration that this concentration correlated with reduction in viremia in bNAb-treated clinical trial subjects (29). The combination of N6 with 10-1074 showed the greatest breadth of coverage, at 83% (IC_80_≤ 10 μg/ml) (**Fig. 2D**), followed by the combination of VRC07-523 and 10-1074, which displayed an IC_80_≤ 10 μg/ml for 81% of the reservoir virus isolates. Several antibody combinations displayed an IC_80_ ≤ 10 μg/ml for 78% of the reservoir virus isolates: N6 and PGT121, VRC07-523 and PGT121, 3BNC117 and 10-1074, 3BNC117 and PG9, 10E8v4-V5R-100cF and 10-1074. Thus, two antibody combinations are able to provide broad neutralization coverage of reactivated reservoir viruses at an IC_80_ ≤ 10 μg/ml for this geographically discrete clade B infected population.

### Infected-Cell Binding Profiles of bNAbs and bNAb Combinations Against Reactivated Reservoir Viruses

We next measured the binding of bNAbs to the surface of primary CD4^+^ T-cells infected with the same reservoir virus isolates that had been assessed for neutralization. Activated CD4^+^ T cells from HIV-uninfected donors were infected with reactivated reservoir viruses and stained with unconjugated bNAbs, followed by Alexa Fluor 647 anti-human IgG secondary antibody. These samples were also stained with HIV Gag to identify infected cells. We used a Median Fluorescence Intensity (MFI) ratio to quantify specific bNAb binding activity to infected cells [MFI ratio = (MFI of bNAb staining in HIV-Gag^+^ cells) / (MFI of bNAb staining in HIV-Gag^-^ cells)] (**Fig 3A**). Since we had already established the geometric mean IC_80_ neutralization values for each virus, we opted to test infected-cell binding at two concentrations for each antibody: i) 5 μg/ml - selected based on titration experiments (data not shown) ii) geometric mean IC_80_ neutralization concentrations for each antibody (values are indicated below the table in **Fig 2C**).

**Figure 3.**
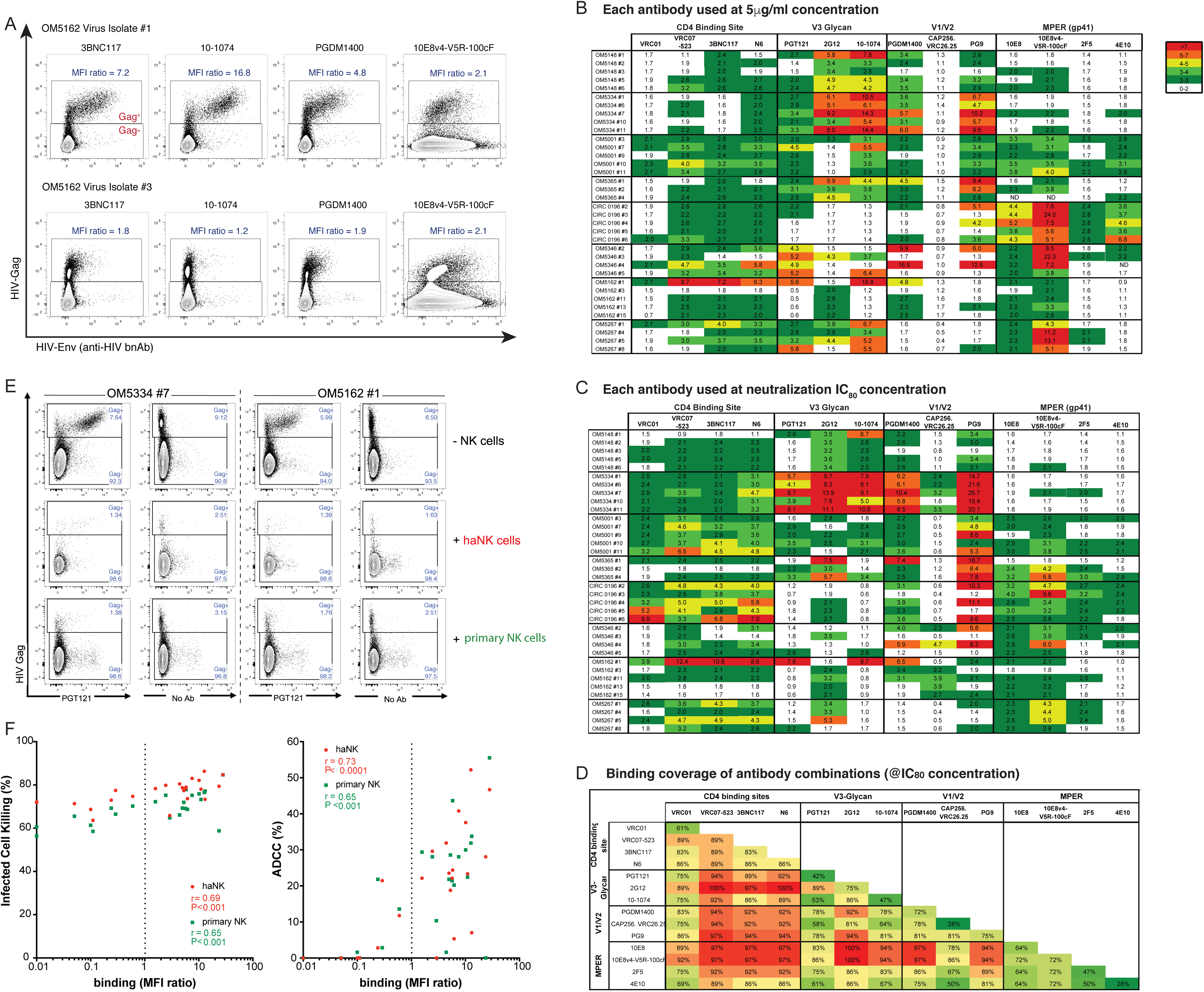
Breadth, potency and functional consequences of binding of a panel of bNAbs against reactivated reservoir viruses. (A) Representative flow plots showing bNAb binding to cells infected with reservoir viruses, gated on live/CD3^+^ cell populations. For each bNAb/virus combination we calculated a median intensity fluorescence (MFI) ratio, defined as MFI of bnAbs in HIV infected cell population (Gag^+^) / MFI of bnAbs in HIV uninfected cell population (Gag^-^). The displayed plots provide an example intra-participant diversity in bNAb binding to different viral isolates. (B) Heat-map showing binding of bNAbs at 5 μg/ml to the indicated viral isolates. The numbers given are MFI ratios, with higher values indicating higher levels of binding. (C) Heat-map showing binding of each bNAb to infected cells when tested at its neutralization geometric mean IC_80_ neutralization concentration. (D) Heat-map showing binding coverage of single bNAbs and two bNAb combinations. The breadth of coverage of antibody combinations was defined based on having at least one of the two bNAbs bind with an MFI ratio > 2. (E) Representative flow cytometry plots from ADCC assays, sampled after the 7 hour co-culture periods and gated on live/CD3^+^ cell populations. The no NK cell conditions (top row) show populations of HIV-infected cells (Gag^+^) that also stain positive for the bNAb PGT121 when added. The addition of either haNK cells (middle row) or primary NK cells (bottom row) resulted in substantial reductions in HIV-infected cell populations, which was generally enhanced by the addition of bNAbs. For the conditions with PGT121, the killing of HIV-infected cells can also be observed in the elimination of cells staining positive for PGT121 in the conditions with NK cells. (F) Correlations between killing frequency (%) and infected cell binding (left panel), and ADCC (%) and infected cell binding (right panel). Both correlations were tested with 2 reservoir viruses combined with 9 bNAbs and the A32 antibody. Each virus/bNAb combination is indicated by a dot, and each color represents one effector cell type, red - haNK cells, green - primary NK cells from the PBMCs of a HIV-negative donor (allogeneic). Correlation coefficients (r) and statistical significance (p) were calculated using Spearman’s Rank-Order Correlation.

In order to establish breadth, we defined binding as a MFI ratio > 2. In general, with the exception of VRC01, CD4bs Abs exhibited superior breadths of infected-cell binding, covering 83 – 89% of reservoir isolates when tested at the neutralization IC_80_ concentrations (**Fig 3C & D**). The binding potencies of CD4bs were relatively modest, however, with most exhibiting MFI ratios of between 2 - 4 (**Fig 3B & C**). The V3-Glycan antibodies PGT121, 2G12, and 10-074 exhibited more limited breadths as compared to CD4bs antibodies, but showed substantially higher levels of specific binding to cells infected with susceptible viruses, with many MFI ratios exceeding 5. Sensitivity/resistance profiles were generally related for different viral isolates from the same individual, e.g. 10-1074 bound strongly to all isolates from 5/8 participants (**Fig 3B**), but exhibited a lack of binding to all viruses from CIRC0196 (at both concentrations). Intra-patient variability was observed, however, for example with 1 out of 5 viruses from OM5162 exhibiting high sensitivity to 10-1074 and the remaining 4 exhibiting resistance. With the exception of CAP256.VRC26.25 [which is predominately clade C specific (22)], the V1/V2 bNAbs showed potent binding activity, particularly in the case of PG9 which, at IC_80_ concentration, showed high levels of specific binding to 16 of 36 reservoir viruses with an MFI ratio greater than 4 (**Fig 3C**). Infected cell binding of MPER-specific antibodies varied: 10E8v4-V5R-100cF (a version of 10E8 optimized for increased solubility and potency(45)) at 5 μg/ml, bound to 30 of 36 isolates, with high-level binding observed for 13 of these (MFI ratios > 4). However, 10E8 and 10E8v4-V5R-100cF also showed substantial binding to uninfected bystanders (Gag^-^ population) (see **Fig 3A**, right panel for representative staining). In contrast, the MPER-specific bNAbs 2F5 and 4E10 exhibited generally narrow and weak binding of reservoir viral isolates (**Fig 3B & C**). Of note, virus #1 from patient OM5162 showed a highly distinct bNAb binding profile as compared to other isolates from the same individual: it was bound strongly by antibodies VRC07-523, 3BNC117, N6, PGT121, 10-1074 and PGDM1400, whereas other autologous viral isolates were bound weakly if at all by these bNAbs. Similarly, viruses from OM5346 showed intra-individual diversity in binding to V3-glycan-specific bNAbs too, as shown PGDM1400 and PG9 bound robustly to viruses #1 and #3 (MFI ratio > 6), while no binding was observed for viruses #2 and #4 (**Fig 3B & C**). Our data indicate both intra- and inter-individual variability in binding to cells infected with reservoir viral isolates, highlighting the limitations of using any single antibody in a therapeutic.

Achieving broad coverage of viral reservoir isolates in a population is likely to require combinations of at least two bNAbs. To assess this in the current population, we calculated the binding coverage of all possible two antibody combinations using the binding data obtained with the neutralization IC_80_ antibody concentration (MFI ratio > 2) (**Fig 3D**). All CD4bs (excluding VRC01) antibodies, when combined with 2G12 or V1/V2 antibodies or MPER antibodies (except for 4E10), reached ≥ 92% coverage. Notably, the combinations of 2G12 with VRC07-523 or N6, or 10E8 or 10E8v4-V5R-100cF reached 100% coverage, however, as previously mentioned, 10E8v4-V5R-100cF showed a high level of bystander binding in our *in vitro* assays. 3BNC117 + 2G12 and VRC07-523 + PG9 reached 97% coverage, thus representing promising combinations for targeting reactivated clade B reservoir viruses (**Fig 3D**).

With respect to the effects of the different concentrations of antibodies tested on binding, 10E8v4-V5R-100cF exhibited generally more favorable binding profiles (MFI ratios) at 5 μg/ml, due to a reduction in the background binding that was observed at its IC_80_ concentration of 9.3 μg/ml. In contrast, 10-1074 showed a lack of background binding even at 5 μg/ml, and thus displayed favorable binding profiles at this higher concentration, compared to its IC_80_ concentration at 0.7μg/ml (**Fig 3B & D**).

### Infected Cell Binding Correlates with Elimination by ADCC

Our primary interest in assessing infected-cell binding is to predict the ability of a bNAb to direct ADCC against these cells. Infected cell binding is a prerequisite for ADCC, and multiple studies have indicated that, where antibody Fc domains are matched (as all bNAbs tested here share the same IgG1), levels of binding correlate with ADCC activity (8, 32-34). To confirm this relationship under our experimental conditions, we performed paired infected cell binding and ADCC assays using two reservoir isolates (OM5334#7 and OM5162#1) in combination with 9 bNAbs. Two types of NK cells were tested in parallel as effectors: i) haNK cells (NantKwest) – a derivative of the NK-92 cell line (46) that has been enhanced for ADCC by expressing high affinity (ha) huCD16 V158 FcγRIIIa receptor, as well as engineered to express IL-2 (47) ii) Freshly isolated NK cells from the peripheral blood of an HIV-uninfected donor. Binding assays were performed in parallel with ADCC assays using the same conditions - 10μg/ml over a total of 7 hours at 37°C. For both haNK cells and primary NK cells, we observed moderate levels of NK-cell mediated elimination of HIV-infected cells in the absence of bNAbs, likely due in part to HIV-mediated downregulation of HLA molecules “missing self” (Fig 3E, F) (48, 49). As expected, we observed additional elimination of infected cells with the addition of bNAbs, and significant direct correlations between total levels of elimination of HIV-infected cells (haNK, r = 0.69, p < 0.001; primary NK cells, r = 0.65, p < 0.001), as well as ADCC-specific elimination of infected cells (% killing in +bNAb conditions - % killing in – bNAb conditions) (haNK, r = 0.73, p < 0.0001; primary NK cells, r = 0.65, p < 0.001) (**Fig 3E, F)**. Thus, our results are consistent with previous studies in indicating that infected cell binding is moderately predictive of ADCC activity for bNAbs with matched Fc domains.

### bNAbs Exert Differential Binding to Populations of Early (Gag^+^CD4^+^) Versus Late (Gag^+^CD4^-^) HIV-Infected Cells

The infection of a cell by HIV results in the progressive, and almost complete, loss of surface CD4 expression, through the concerted actions of Nef, Vpu, and Env(50-53). Thus, in short-term *in vitro* infections of activated CD4^+^ T-cells, Gag^+^CD4^-^ cells represent a later stage of infection than their Gag^+^CD4^+^ counterparts (which have not yet downregulated CD4). Env is expressed at substantially higher levels in late-versus early-infected cells. Thus, variations in overall levels of antibody binding to total Gag^+^ cells between different viral isolates, as observed in **Fig 3**, may reflect not only intrinsic differences in bNAb sensitivity but also differences in infection kinetics (different ratios of early: late infected cells and therefore different levels of protein expression). We therefore sought to refine our analysis of levels of bNAb binding by controlling for stage of infection.

We assessed whether differential binding of bNAbs to early (Gag^+^CD4^+^) versus late (Gag^+^CD4^-^) infected cells was present in our assays. Upon gating on viable HIV-infected cells (lymphocytes, live cells, CD3^+^, HIV-Gag^+^), we observed that some bNAbs, such as 3BNC117, 10-1074, and PG9 showing preferential binding to late infected cells (CD4^-^) (**Fig 4A**), while others, such as 10E8v4-V5R-100cF, showing similar, or slightly higher binding to early versus late populations (**Fig 4A**). To test this systematically, we selected virus/bNAb combinations that showed specific binding (Gag^+^/ Gag^-^ bNAb MFI ratio > 2 when tested at neutralization IC_80_ concentrations) and compared levels of bNAb binding in the Gag^+^CD4^+^ versus Gag^+^CD4^-^ populations. We calculated fold differences between these early and late infected populations = (Geometric Mean MFI ratio of Gag^+^CD4^-^) / (Geometric Mean MFI ratio of Gag^+^CD4^+^). We observed that all gp120-specific bNAbs exhibited higher levels of binding to late-infected populations than to matched early-infected populations (Fold differences: 1.7 – 3.9, **Fig 4B**). In contrast, each of the gp41-specific bNAbs exhibited similar or slightly higher levels of binding to early-versus late-infected populations (Fold differences: 0.90 – 0.97, **Fig 4B**). A mechanistic explanation for this discrepancy is beyond the scope of the current manuscript. However, we raise the possibility that it may be related to the in *cis* interactions that have been shown to occur on early infected cells between gp120 and CD4 on the same cell surface(54). Binding of CD4 to functional trimers can induce gp120 shedding from Env trimers, and enhance exposure of the gp41 membrane proximal external region to antibody binding (55). Our data are consistent with such conformational differences in Env favoring gp41-specific antibody binding to early-infected cells.

**Figure 4.**
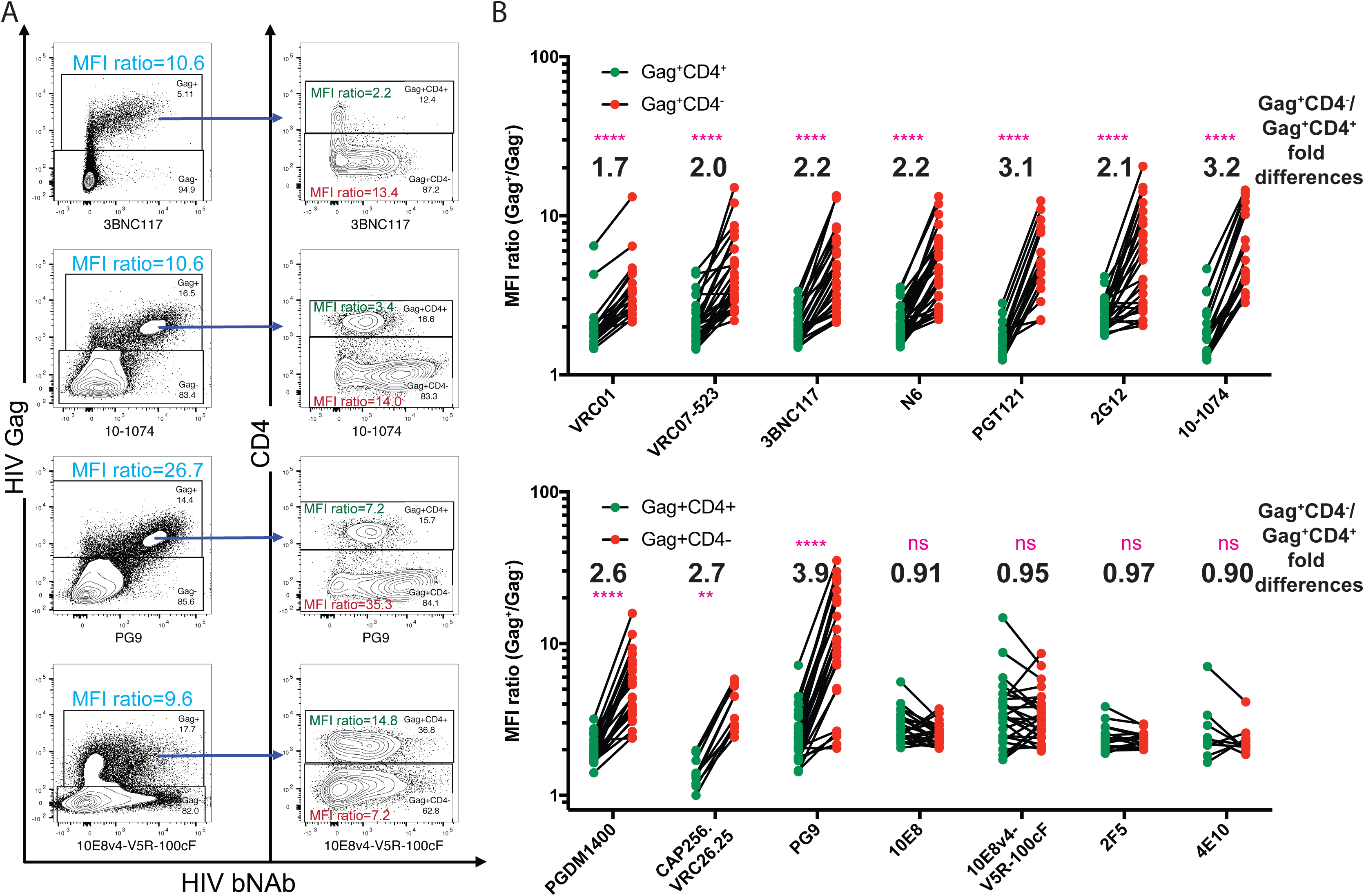
Comparisons between bNAb binding to early (Gag^+^CD4^+^) versus late (Gag^+^CD4^-^) HIV-infected cell populations. Gag^+^CD4^-^ population represents the specific binding to HIV Env. (A) Representative flow cytometry plots gated on lymphocytes/live/CD3^+^ (left panels), and on lymphocytes/live/CD3^+^/HIV-Gag^+^ (right panels) showing differential bNAb binding to CD4^+^ (early infected) and CD4^-^ (late infected) populations. The results show that for 3BNC117, 10-1074 and PG9 most of the bNAb binding to infected cells (Gag^+^) occurs with the CD4^-^ population. In contrast, for 10E8v4-V5R-100cF CD4^+^ T-cells are bound at similar or slightly higher levels than CD4^-^ T-cells (within the Gag^+^ population). (B) Summary data for the analysis represented in panel A, showing paired comparisons of MFI ratios between CD4^+^ and CD4^-^ populations. MFI ratio is defined as (MFI of bNAbs in Gag^+^CD4^+^)/ (MFI of bNAbs in Gag^-^) (green dots) or (MFI of bNAbs in Gag^+^CD4^-^)/ (MFI of bNAbs in Gag^-^) (red dots). The numbers indicate fold differences (mean of Gag^+^CD4^-^ MFI ratio)/ (mean of Gag^+^CD4^+^ MFI ratio) (Wilcoxon matched-pairs signed rank test, **** p<0.0001, ** p<0.01).

One implication of these results is that the binding data presented in **Fig 3** - which was generated based on total Gag^+^ cells – over-represents binding to early-infected cells for gp120-specific antibodies, and under-represents binding to late-infected cells. Data calculated based only on the late-infected populations show a substantially intensified binding profile for most of the bNAbs used in this study – most notably for the CD4bs bNAbs and PG9 (binding @ geographic mean IC_80_ concentration, **Supplementary Fig 1**). A second implication is that cellular infection dynamics may impact the ability to detect relationships between infected-cell binding and virus neutralization. For example, if virus 1 replicated with faster kinetics than virus 2, and thus had a greater proportion of Gag^+^CD4^-^ versus Gag^+^CD4^+^ cells, then this would skew bNAb binding profiles in a way that was not intrinsic to the Env itself. To account for this, we have assessed these relationships based on both total Gag^+^ cells and on only the Gag^+^CD4^-^ late infected populations (below).

### Virus Neutralization Correlates with Infected-Cell Binding for most bNAbs

The breadths and potencies of neutralizing activity of bNAbs against diverse HIV isolates have been extensively studied (13-22). In contrast, relatively few studies have assessed breadths and potencies of infected-cell binding, which is an important pre-requisite for ADCC (8, 32-34). Efforts to harness bNAbs to direct ADCC against infected cells would therefore benefit from an understanding of the degree to which infected cell binding can be inferred from neutralizing activity against a given virus. Our paired binding and neutralization data sets allowed us to assess this using a number of analytic approaches in regards to both concentrations of bNAbs used for binding assays and to stage of infection of target cells. With respect to bNAb concentrations, binding to infected cells was assessed for each bNAb at 5 μg/ml, and at the geometric mean IC_80_ neutralization concentration of that antibody against the same panel of reservoir viruses. For the latter, this meant that some antibodies were tested at > 5 μg/ml (ex. 4E10 at 49.2 μg/ml), while other antibodies were tested at substantially lower concentrations (e.g. PGT121 at 0.6 μg/ml) (Geo Mean IC_80_ concentrations are given below the heat-map in **Fig 2C**). This approach thus seeks to normalize for intrinsic differences in avidity between different bNAbs. With respect to stage of infection of target cells, we separately tested for correlations between neutralization IC_80_ and binding to either total infected cells (Gag^+^) or to late-infected cells (Gag^+^CD4^-^), based on the differential binding patterns described above. Of these, the most appropriate method for assessing the relationship between binding and neutralization likely depends on the question being asked. Importantly, however, the relationships that we observed, as described below, turned out to be conserved across these different approaches.

We first tested for correlations between neutralization IC_80_ and the level of binding (MFI ratio) at 5 μg/ml bNAb concentrations. As is described above, since cells in early versus late stages of HIV infection exhibit differential bNAb binding profiles, replication dynamics have the potential to impact overall assessments of binding. In order to increase our ability to discern Env-intrinsic relationships between binding and neutralization we therefore limited this initial analysis to the late-infected (Gag^+^CD4^-^) population. When all antibodies were considered together, we observed a significant, direct correlation between virus neutralization and infected cell binding (p < 0.0001, Spearman’s r = 0.63) (**Fig 5A**). For each of the bNAbs that showed appreciable neutralizing activity (VRC01, VRC07, 3BNC117, N6, PGT121, 10-1074, PGDM1400, PG9, 10E8, and 10E8v4-V5R-100cF) we observed significant direct correlations between neutralizing activity and infected-cell binding (**Fig 5B)**. The antibodies 2F5 and CAP256.VRC26.25 showed little in the way of either neutralization or binding, precluding the possibility of detecting a relationship between these factors. 2G12 and, to lesser extent, 4E10 were notable outliers as they showed appreciable binding capacity to many of the viruses in this panel, but very little corresponding neutralizing activity. This lack of potent neutralization activity is inconsistent with data from pseudovirus assays, but in agreement with previous data using virus produced from T-cells, suggesting that 2G12 sensitivity is particularly tied to the source of virus (56-58).

**Figure 5.**
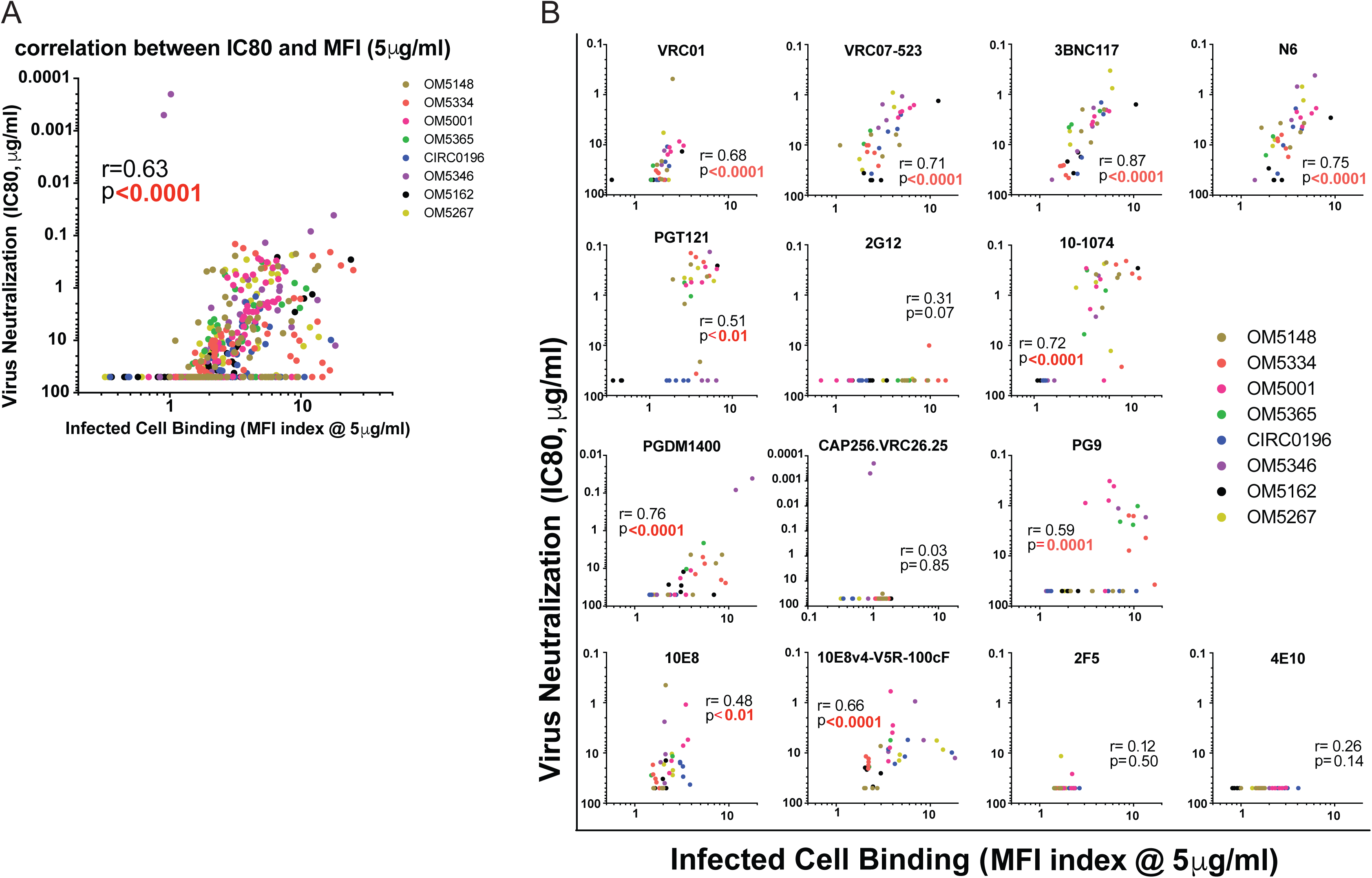
Correlations between virus neutralization and paired late-infected cell binding at 5 μg/ml bNAb concentrations. Shown are correlations between IC_80_ virus neutralization values and binding to late-infected cells (Gag^+^CD4^-^) using a 5 μg/ml concentration for each antibody. (A) Correlation for all antibodies tested together. (B) Correlations for each bNAb tested independently. Each virus/bNAb combination is indicated by a circle, and each color represents one study participant. Correlations were analyzed by Spearman correlation coefficient (r), with statistical significance highlighted in red letters.

Correlations between neutralization IC_80_ and binding as measured by other approaches are presented as follows: i) binding of antibodies tested at 5 μg/ml concentrations to total infected population (all Gag^+^) – **Supplementary Fig 2;** ii) binding of antibodies tested at IC_80_ neutralization concentrations to late infected population (Gag^+^CD4^-^) – **Supplementary Fig 3;** iii) binding of antibodies tested at IC_80_ neutralization concentrations to total infected population (all Gag^+^) – **Supplementary Fig 4**. Correlation coefficients varied across these different analyses, with different approaches yielding stronger correlations for different bNAbs, e.g. for 3BNC117: Spearman’s r = 0.82 for 5 μg/ml total Gag^+^ (**Supplementary Fig 2**) vs Spearman’s r = 0.60 for IC_80_ concentration total Gag^+^ (**Supplementary Fig 4**); for PGT121: Spearman’s r = 0.47 for 5 μg/ml total Gag^+^ (**Supplementary Fig 2**) vs Spearman’s r = 0.71 for IC_80_ concentration total Gag^+^(**Supplementary Fig 4**). Overall, however, each of the antibodies that exhibited a significant correlation by one analytic approach also exhibited significant correlations by the other three approaches, and vice versa for those lacking significant correlations. Thus, for 10 out of 14 bNAbs tested in this study, the ability of a bNAb to neutralize a given virus is strongly correlated with its ability to bind to a corresponding infected cell. In these *in vitro* assays, this correlation was robust enough to be observed with or without controlling for avidity of a given bNAb or for infection dynamics.

## Discussion

The primary conclusion of the current study is that the ability of a given bNAb to neutralize clinical viral isolates is a strong correlate of its ability to bind to cell-surface Env on primary CD4^+^ T-cells infected with the same virus. Furthermore, in comparing across a large panel of bNAbs, relative levels of infected-cell binding and virus neutralization continued to correlate – for example, 10-1074 showed both high-level infected-cell binding and potent neutralization compared to VRC01. Thus, we conclude that – with respect to the Fab component of Abs, when sharing the same Fc – the selection of Abs based on broad and potent neutralizing activity is very likely to also select for those that are suitable for infected-cell clearance. Of note, the reciprocal was not always true; with 2G12 exhibiting reasonably potent and broad infected-cell binding, contrasted by a general lack of neutralization of these reservoir-derived primary isolate viruses. Though less strikingly, the MPER-specific bNAbs 2F5 and 4E10 also exhibited appreciable infected-cell binding (similar in breadths and magnitudes to VRC01), but with minimal neutralizing activity. We propose that the differences based on the directionality of this relationship may be related to the differential antigen conformational requirements for these two functions. For a bNAb to neutralize virus, it must bind functional Env trimers present on the surface of cells producing infectious virus. In contrast, an antibody that also binds to nonfunctional envelope proteins, such as gp41 stumps (59), may bind to infected cells to a greater degree than they mediate neutralization (if they neutralize at all). Thus, virus neutralization is a predictor of infected-cell binding, but the reciprocal relationship does not hold.

While it may be intuitive that virus neutralization would correlate with infected-cell binding, we do not feel that this could have been assumed to be the case without experimental evidence. The conformation of Envs may be affected by differences between the cell-surface vs virion environments, and this variability could impact different viral isolates. For example, in *cis* interactions between CD4 and Env on the surfaces of infected cells have been shown to induce gp120 shedding, and expose gp41 stumps. This has been reported to enhance infected-cell binding by gp41-specific Abs, while diminishing binding by gp120-specific Abs (35). Such an effect might differentially impact different viruses – for example, Horwitz *et al*. reported that the R456K mutation on YU2 gp120 decreased gp120 shedding, which led to less bystander (Gag^-^CD4^+^) binding (60). Our data are consistent with these observations, and provide further evidence of *cis* binding of CD4 modulating the binding of bNAbs to infected-cells. We find gp120-specific bNAbs bind preferentially to cells in a late stage of infection (CD4^low^) while gp41-specific bNAbs bind similarly or slightly better to cells in an early stage of infection (CD4^high^). To address more mechanisms of these findings, future studies may benefit from including gp120/gp41 interface bNAbs, such as 8ANC195 (61), PGT151, PGT158 (62). However, despite any such differences between the virion and cell-surface environments, the ability to neutralize virus was significantly correlated with infected-cell binding, and these relationships held whether we considered all infected cells (Gag^+^) or only late infected cells (Gag^+^CD4^-^).

To investigate factors that may predict the efficacy of bNAb treatment to contribute to HIV cure we felt it important to study the properties of bNAbs against viruses derived from reactivated latent reservoirs. By combining a QVOA approach with isolation of virus from dilutions of CD4^+^ T cells from different ART-suppressed patients where <50% of wells were p24^+^, we were able to isolate viruses that were likely clonal to test bNAb binding and neutralization profiles (**Fig 1**) and assess both intra-and inter-patient variability. We observed a considerable level of heterogeneity, even within a given individual, such that in the majority of cases any single bNAb failed to provide universal coverage of an individual’s reservoir isolates. However, combinations of two antibodies provided broad coverage both within and across individuals, reaching up to 100% coverage as assessed by binding. Note that as our study population was derived from a single site (Toronto, Canada), from a clinical perspective this assessment of breadth is representative of what might be expected in a single-site study in a North American clade B infected cohort. We propose that the method presented here could be applied to different populations as a means of prioritizing antibody combinations for a given regional population of patients and personalizing individual HIV cure strategies as ART drug resistance is used to guide ART therapy. Clinical use of the QVOA assay will likely be limited by its expense, cell number requirements, and protracted timeline (14 days) for results. However, a notable opportunity is present in the fact that infectious clonal autologous reservoir viruses are generated as a byproduct of the primary measurement. The pairing of <u>q</u>uantitative and <u>q</u>ualitative assessments of the HIV reservoir in this way has been previously termed the Q^2^VOA (63).

The potencies of neutralization observed in the current study are overall weaker than those that have been previously reported using pseudovirus assays – most notably for 2G12, which failed to achieve 80% neutralization for all but two viruses. While this is likely due in part to our use of clinical viral isolates, which are generally less sensitive to bNAbs than laboratory-adapted viruses (64, 65), we also note the role of virus producing cells in modulating sensitivity to neutralization. Studies addressing this issue have reported that T-cell derived virus is more resistant to neutralization than pseudovirus generated by transfected 293T cells and, in particular, that replication competent virus produced by PBMCs are more neutralization resistant than Env matched pseudoviruses (56-58). However, there appear to be antibody-specific differences in the level of influence that a producer cell has on sensitivity to neutralization. For example, one study reported that PG9 is not very sensitive to differences in producer cell (66), while large differences in IC_50_ have been reported between T cell and pseudovirus for antibody 2G12 (56, 57). These data suggest that producer cells differentially influence the conformations of Env on resulting virions, as well as their densities and glycosylation, or numbers of gp120 molecules in the viral membrane. As PG9 preferentially targets well-ordered, closed, trimeric viral spikes, it indicates that an equal number of well-folded spikes exists on virions produced by either cell type, whereas perhaps bNAbs such as 2G12 can bind equally well to mis-folded trimers and are therefore more sensitive to increases in the latter. Furthermore, the epitopes of certain antibodies, such as 2G12, include glycans, and producer cells can affect glycosylation patterns of gp120 (66). Thus, in addition to the comparison between neutralization and infected-cell binding, the current study contributes a reassessment of bNAb neutralization potency that may be more clinically applicable than data from pseudovirus assays.

In conclusion, our study provides novel insights into the relationship between infected-cell binding and virus neutralization that may help to guide immunotherapeutic strategies aimed at either curing infection, or enabling durable immune control of viral replication. The degree of intra-and inter-individual variation in bNAb sensitivity within even this geographically discrete clade B population reinforces the importance of utilizing combinations of at least two bNAbs in such therapies. Screening reactivated reservoir viruses for sensitivity to bNAbs, either at an individual or population level, can help select antibody combinations for optimal coverage – for example, with combinations of PG9 and either 3BNC117 or N6 providing potent infected-cell binding coverage of 94% and 72-78% coverage of neutralization (IC_80_ ≤ 10μg/ml) of viruses in the current study population. For the bNAbs that exhibited correlations between infected-cell binding and neutralization, our study indicates that screening for either one of these factors is sufficient to infer that both functions will be present against reactivated reservoir viruses. Consistent with previous studies, we also confirmed that this infected cell binding – as measured by our assay – correlated well with NK cell mediated ADCC, suggesting that it is a reasonable surrogate. It will be of interest, however, for future studies to build upon these results with more extensive functional assays (potentially using varying Fc domains and/or effector cells). Such future directions could potentially uncover more subtle aspects of the relationship between virus neutralization and the targeting of cell-mediated Fc-dependent functional activities against infected cells, which may lead to the elimination of latent reservoirs.

## Materials and Methods

### Ethics Statement

All participants (HIV-infected individuals) were recruited from the Maple Leaf Medical Clinic in Toronto, Canada, through a protocol approved by the University of Toronto Institutional Review Board. Secondary use of the samples from Toronto was approved through the George Washington University Institutional Review Boards. All subjects were adults, and gave written informed consent. Clinical data for these participants are given in **Table 1**.

### Broadly Neutralizing Antibodies

We used a panel of broadly neutralizing antibodies to HIV (bNAbs): CD4 binding sites antibodies (CD4bs)-VRC01, VRC07-523, 3BNC117, N6; V3-Glycan antibodies-PGT121, 2G12, 10-1074; V1/V2 antibodies-PGDM1400, CAP256.VRC26.25, PG9; MPER antibodies-10E8, 10E8v4-V5R-100cF, 2F5, 4E10; and a positive control antibody HIV-IG and a negative control antibody 4G2-Hu for neutralization assays. Antibodies 10-1074, 2G12, and control antibody HIV-IG were obtained through the AIDS Reagent Program, Division of AIDS, NIAID, NIH from Dr. Michel C. Nussenzweig, Polymun Scientific and NABI and NHLBI, respectively. Dr. John Mascola provided antibody proteins 2F5 and 4E10, as well as all other antibody heavy-and light-chain expression plasmids. Antibody plasmids were expressed as full-length IgG1s from transient transfection of 293F cells and purified by affinity chromatography using HiTrap Protein A HP Columns (GE Healthcare).

### Quantitative viral outgrowth assay (QVOA)

Human CD4 T cells were enriched from the peripheral blood mononuclear cells (PBMCs) (Stemcell Technologies), processed from leukapheresis, which were drawn from long-term ARV-treated HIV-infected participants (**Table 1**). Cells were diluted into a serial concentration (2 million, 1 million, 0.5 million, 0.2 million and 0.1 million per well), and plated out into 24 well-plates and each concentration would have 12 wells. PHA and irradiated PBMCs were added to reactivate the infected cells and MOLT-4 cells were added 24 hours later to amplify the viruses. Media were changed every 3-4 days and p24 ELISA were run on day 14 to measure the amount of virus production.

### p24 Enzyme-Linked Immunosorbent Assay

p24 enzyme-linked immunosorbent assay (ELISA) was performed with kit components obtained from National Cancer Institute, NIH. In brief, 96-well high binding microplates (Greiner Bio-One) were coated with capture antibody for overnight, and followed by 1% BSA solution blocking for overnight. Supernatants from QVOA wells were collected and lysed with 1% x-Triton buffer for 2 hours, followed by transferring to ELISA plates and incubating for 1 hour, 37°C. Plates were then washed with PBST buffer (PBS+0.1% Tween-20) for 6 times and incubated with primary antibody for 1 hour, 37°C. After 6 additional washes, peroxidase labeled Goat anti-rabbit IgG secondary antibody (KPL) was added and incubated for another 1 hour at 37°C. After 6 additional washes, TMB substrate (Thermo Fisher) was added and developed for 15 mins, then stopped with stop solution (Biolegend). Absorbance was measured with SpectraMax i3x Multi-Mode microplate reader (Molecular Device) at OD450nm and 570nm. Cut offs for positive wells were set as > 2x the average of negative control values.

### Neutralization assay

Neutralization of QVOA virus samples by bNAbs were measured using infection of Tzm-bl cells as described previously(30, 67). p24 protein in each virus sample was quantified by using the AlphaLISA HIV p24 Biotin-Free detection kit (Perkin Elmer, Waltham, MA), and input virus was normalized to 5-10ng/ml for the assay. 10μl of five-fold serially diluted mAbs from a starting concentration of 50μg/ml were incubated with 40μl of replication competent virus samples in duplicate for 30 minutes at 37°C in 96-well clear flat-bottom black culture plates (Greiner Bio-One). Tzm-bl cells were added at a concentration of 10,000 cells per 20μl to each well in DMEM containing 75μg/ml DEAE-dextran and 1μM Indinavir. Cell only and virus only controls were included on each plate. Plates were incubated for 24 hours at 37°C in a 5% CO_2_ incubator, after which the volume of culture medium was adjusted to 200μl by adding complete DMEM containing Indinavir. 48 hours post-infection, 100μl was removed from each well and 100μl of SpectraMax Glo Steady-Luc reporter assay (Molecular Devices, LLC., CA) reagent was added to the cells. After a 10-min incubation at room temperature to allow cell lysis, the luminescence intensity was measured using a SpectraMax i3x multi-mode detection platform per the manufacturers’ instructions. Neutralization curves were calculated comparing luciferase units to virus-only control after background subtraction and fit by nonlinear regression using the assymetric five-parameter logistic equation in GraphPad Prism (**Fig 2A**). The 50% and 80% inhibitory concentrations (IC_50_ and IC_80_, respectively) were defined as the antibody dilution that caused a 50% and 80% reduction in neutralization.

### bNAb binding assay

All binding assays were tested with the unconjugated bNAbs. CD4^+^ T cells (which were all CD3^+^) were isolated with the Human CD4 T cell enrichment kit (Stemcell Technologies) and activated with CD3/28 antibodies (Biolegend) for 48 hours. Supernatants collected from QVOA wells (p24^+^, the same viruses with neutralization assay) were used for infection by adding into the activated CD4^+^ T cells, followed by spinnoculation for 1 hour and 6 days in culture with media change every 3 days. Infection rate was checked on days 3 and 5 post infection. When most of the infection reached >5%, bNAb staining were performed. Cells were collected and washed twice with 2% FBS PBS, and then aliquoted into 96-well plates (1 million cells per well). Unconjugated bNAbs were added according to the outlined wells by diluting to a final concentration of 5μg/ml or neutralization IC_80_ concentration, which was the Geo Mean of neutralized virus that generated from neutralizing assay, and then incubated at 37°C for 1 hour. Without washing, the Alexa Fluor 647 labeled secondary antibody (Southern Biotech) was added and incubated at 4°C for 30 minutes. After washing once with 2% FBS PBS, surface antibodies mixture was added: BV786 anti-human CD3 (SK7, BD Biosciences), Pacific Blue anti-human CD4 (RPA-T4, BD Pharmingen) and LIVE/DEAD aqua (Life technology). 30 minutes later, cells were washed twice and fixed/permeabilized with Fixation/Permeabilization Solution (BD Bioscience). Anti-HIV-1 core antigen antibody (KC57-RD1, Beckman Coulter) was used to stain intracellular HIV-1 gag protein. After two washes with 1x Perm/Wash buffer, cells were detected by flow cytometry (BD Fortessa X-20), and data analysis was performed with flowjo v10 (Treestar).

### Antibody mediated NK cell killing (ADCC) assay

ADCC assays were performed with unconjugated bNAbs and one of two types of NK cells: haNK cells (NantKwest), a NK-92 cell line which has been engineered to express the high affinity (ha) CD16 V158 FcγRIIIa receptor, as well as engineered to express IL-2 (47); and primary NK cells enriched from the PBMCs of an HIV-negative donor (buffy coat from Gulf Coast Regional Blood Center) using the Human NK cell enrichment kit (Stemcell Technologies). To generate target cells, primary CD4^+^ T-cells were enriched from the PBMCs of allogeneic healthy donors and infected with reservoir viruses as for binding assays (see above). Infections were monitored by flow cytometry, and ADCC assays were performed when target cells were >5 % infected. Both types of NK cells were treated with 10nM IL-15 superagonist complex, ALT-803 (68, 69) for 1 hour to prime and activate them. Infected cells were collected and washed twice with 2% FBS PBS. 2×10^5^ cells/well were added into U-bottom 96-well plates. Unconjugated bNAbs (VRC01, VRC07-523, 3BNC117, N6, PGT121, 2G12, 10-1074, PGDM1400, PG9, A32 or no Ab) were added to final concentrations of 10μg/ml, and then incubated at 37°C for 2 hours. After this incubation, 4×10^5^ ALT-803 treated NK cells were added to each well to give E:T ratios of 2:1. bNAbs binding assays was performed in parallel with the ADCC assay with same conditions but no NK cells added. Plates were centrifuged at 100×g for 30 seconds to bring target and effector cells into contact with each other, and then incubated at 37°C, 5% CO_2_. Cells were mixed by pipetting after 2 hours of incubation, and then cocultured for an additional 5 hours. After a total of 7 hours of co-culture, cells were washed twice with 2% FBS PBS, and stained with fluorochrome-conjugated antibodies against: human IgG, CD3, CD56, CD4 (all from Biolegend), as well as with a live/dead aqua amine reactive dye (Molecular Probes). Cells were then fixed and permeabilized using the BD cytofix/cytoperm kit and following the manufacturer’s instructions. Intracellular HIV-Gag was then stained with PE-conjugated anti-HIV-Gag (clone KC57, Beckman Coulter). Cells were analyzed by flow cytometry (BD Fortessa X-20), and data analysis was performed using flowjo v10 (Treestar). Frequencies of viable Gag^+^ cells amongst the CD3^+^ cells (all targets) were determined. Killing (%) values were calculated using the following formula: [% Gag^+^ (of viable CD3^+^ cells) in no NK cell no Ab condition - % Gag^+^ (of viable CD3^+^ cells) in test condition] / [% Gag^+^ (of viable CD3^+^ cells) cells in no NK cell no Ab condition] * 100%. ADCC (%) values were calculated using the following formula: [% Gag^+^ (of viable CD3^+^ cells) in +NK cells but no Ab condition - % of Gag^+^ (of viable CD3^+^ cells in test condition) / (% of Gag^+^CD3^+^ cells +NK but no Ab condition) * 100%. Negative values were set equal to zero.

### Statistical analysis

Statistical analyses were performed using Prism 7 (GraphPad). Flow data were analized with flowjo v10. The heat-maps were generated with Excel. Comparison between MFI ratio of Gag^+^CD4^+^ and that of Gag^+^CD4^-^ was using Wilcoxon matched-pairs signed rank test. All correlations were calculated using using Spearman’s Rank-Order test.

## Acknowledgments

We thank all of the study participants who devoted time to our research. We also thank Kiera Clayton for helpful comments on the manuscript. Research reported in this publication was supported by the National Institute of Allergy and Infectious Diseases of the National Institutes of Health under award number UM1AI126617 – the Martin Delaney ‘BELIEVE’ Collaboratory, with co-funding support from the National Institute on Drug Abuse, the National Institute of Mental Health, and the National Institute of Neurological Disorders and Stroke. This work was also supported under NIH award numbers AI22391, AI31798, MH12224, and by the NIH funded Center for AIDS Research grant P30 AI117970 which is supported by the following NIH Co-Funding and Participating Institutes and Centers: NIAID, NCI, NICHD, NHLBI, NIDA, NIMH, NIA, FIC, and OAR. The content is solely the responsibility of the authors and does not necessarily represent the official views of the National Institutes of Health. The following materials were supplied by the NIH AIDS Research and Reference Reagent Program: broadly neutralizing antibodies, IL-2, MOLT-4 CCR5 cells.

## Supplementary figure legends

**Fig S1.**
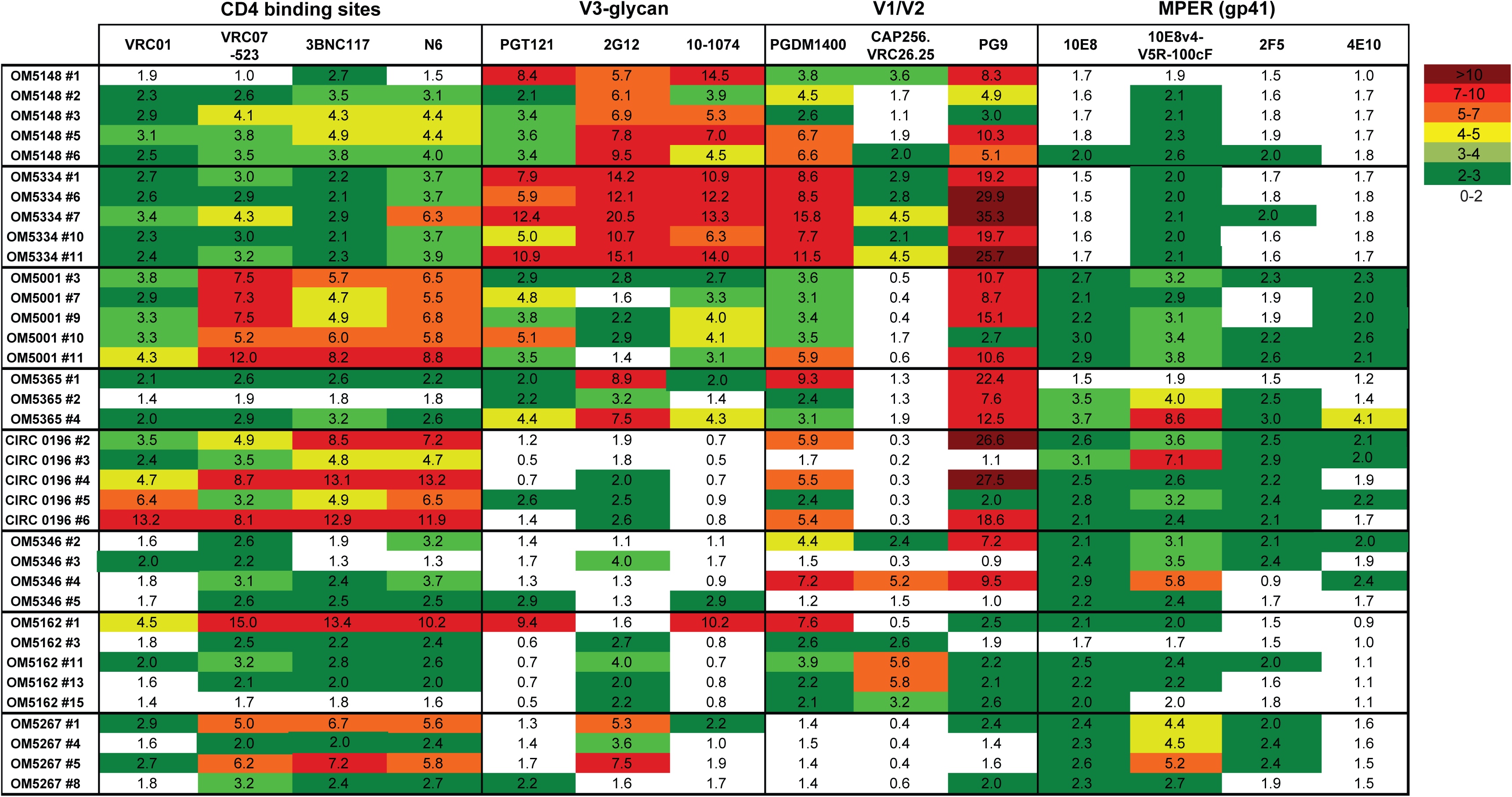
Heat-map of bNAb binding to late-infected (Gag^+^CD4^-^) cell populations at geometric mean IC80 neutralization concentrations. Binding assay were performed with individual geometric mean neutralization IC_80_ concentrations for each bNAb. The numbers indicate MFI ratios of the Gag^+^CD4^-^ population (late infected) / Gag^-^ population (uninfected). Thus a higher value represents a higher level of specific bNAb binding.

**Fig S2.**
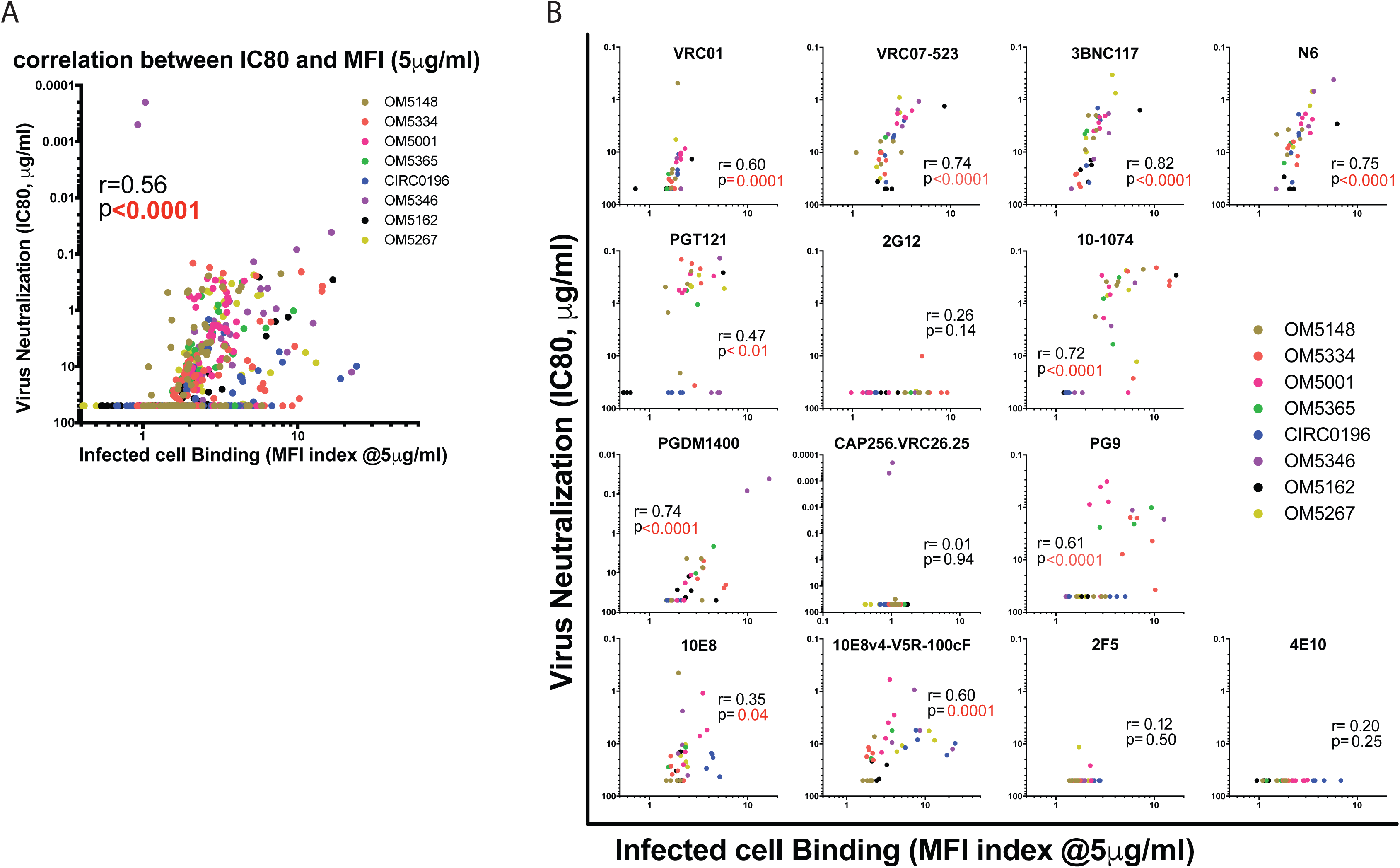
Correlations between virus neutralization and bNAb binding to total infected population (all Gag^+^) when tested at 5 μg/ml. Shown are correlations between IC_80_ virus neutralization values and binding to HIV-infected cells (total Gag^+^, thus groups early and late infection) with each bNAb tested at 5 μg/ml (A) Correlation for all antibodies tested together. (B) Correlations for each bNAb tested independently. Each virus/bNAb combination is indicated by a circle, and each color represents one study participant. Correlations were analyzed by Spearman correlation coefficient (r), with statistical significance highlighted in red lettering.

**Fig S3.**
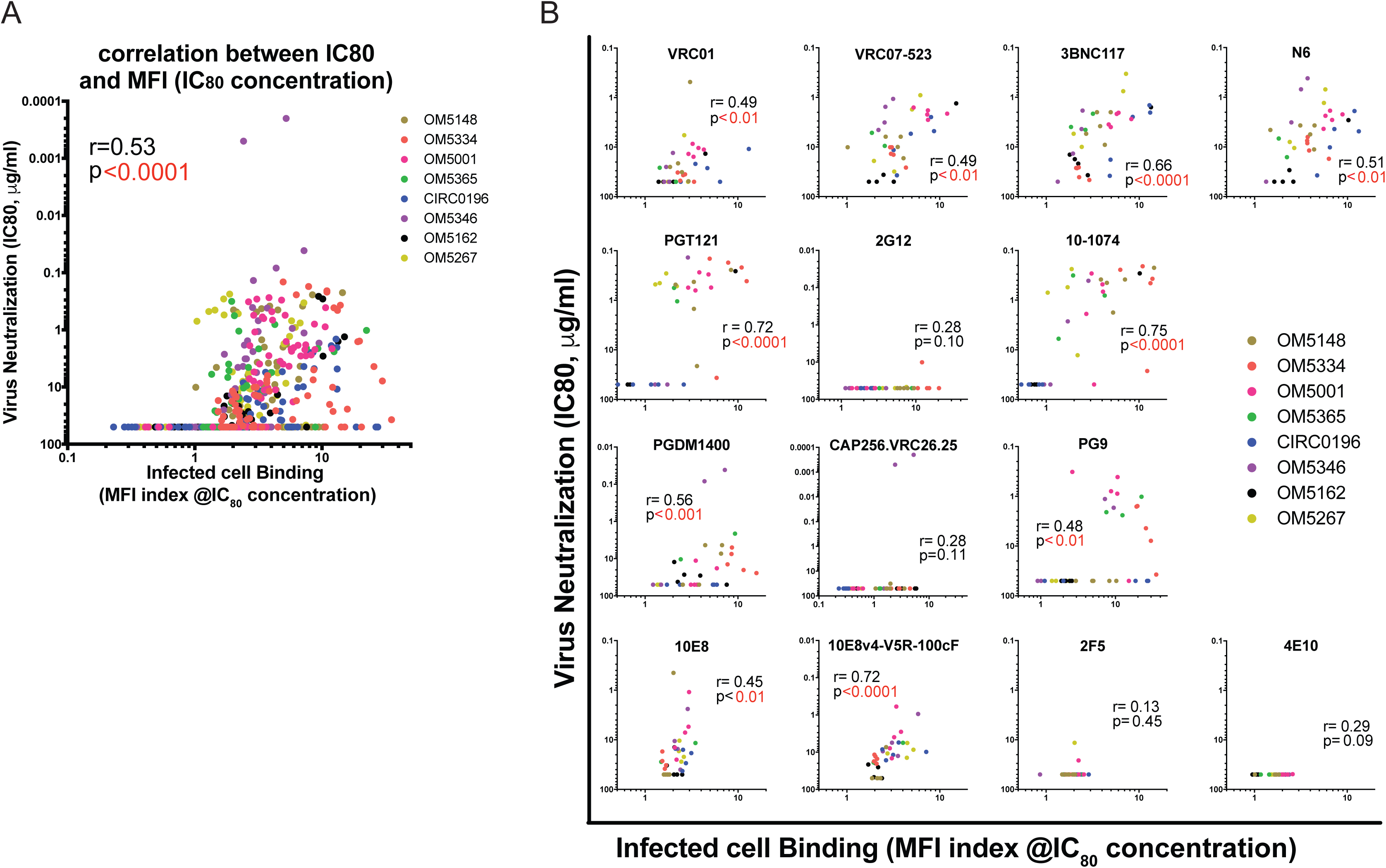
Correlations between virus neutralization and bNAb binding to late-infected populations (all Gag^+^/CD4-) when tested at neutralization geometric mean IC80 concentrations. Shown are correlations between IC_80_ virus neutralization values and binding to late-infected cells (Gag^+/^CD4^-^) with each bNAb tested at its individual geometric mean neutralization IC_80_ concentration (A) Correlation for all antibodies tested together. (B) Correlations for each bNAb tested independently. Each virus/bNAb combination is indicated by a circle, and each color represents one study participant. Correlations were analyzed by Spearman correlation coefficient (r), with statistical significance highlighted in red lettering.

**Fig S4.**
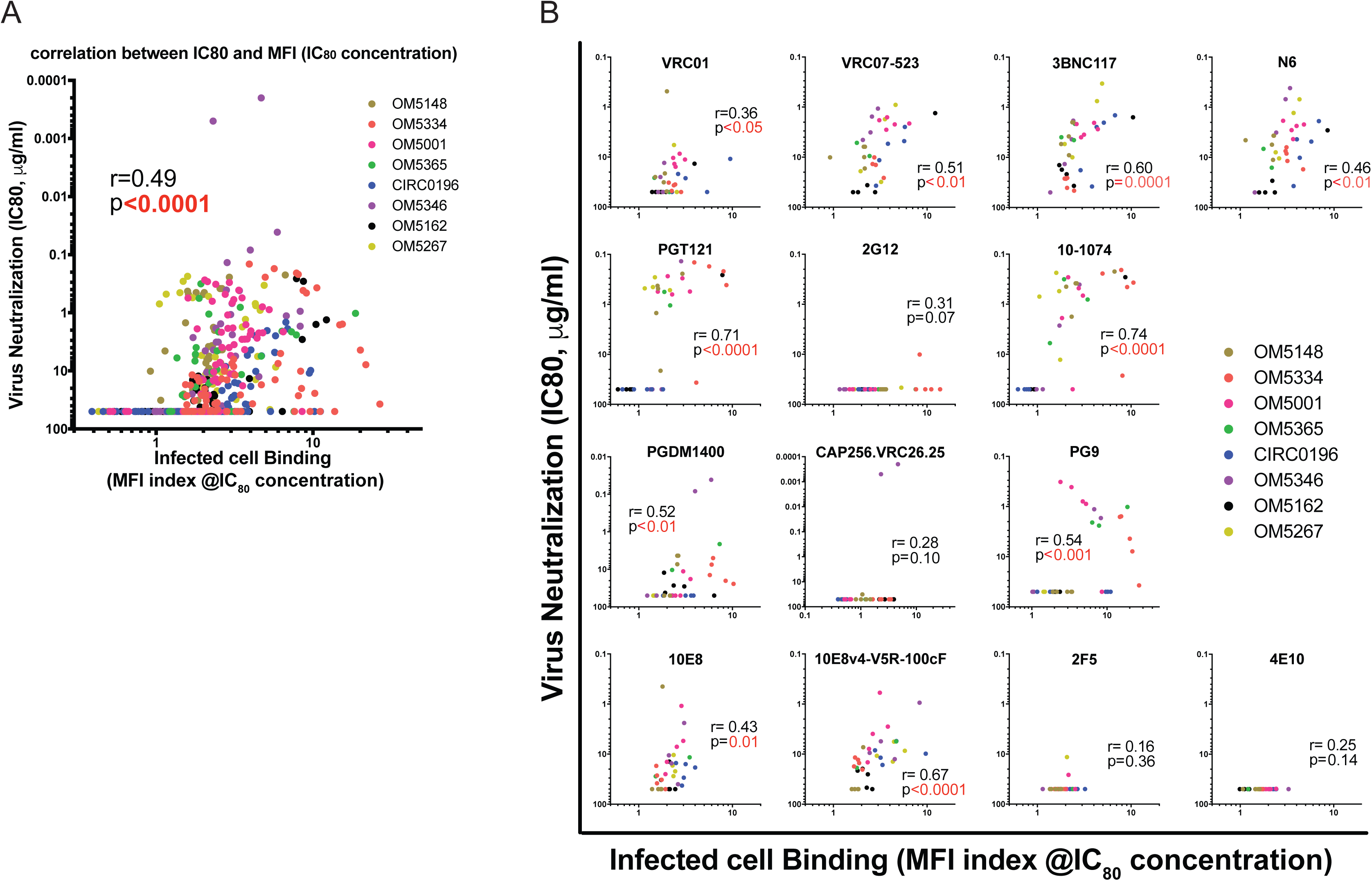
Correlations between virus neutralization and bNAb binding to total infected population (all Gag^+^) when tested at geometric mean neutralization IC80 concentrations. Shown are correlations between IC_80_ virus neutralization values and binding to HIV-infected cells (total Gag^+^, thus groups early and late infection) with each bNAb tested at individual neutralization geometric mean IC_80_ concentrations (A) Correlation for all antibodies tested together. (B) Correlations for each bNAb tested independently. Each virus/bNAb combination is indicated by a circle, and each color represents one study participant. Correlations were analyzed by Spearman correlation coefficient (r), with statistical significance highlighted in red lettering.

